# Epigallocatechin gallate and thermal cycling-stimulation synergistically promote apoptosis in A549 cells via endoplasmic reticulum stress-induced calcium ion dysregulation and oxidative stress

**DOI:** 10.64898/2026.06.23.733932

**Authors:** Fang-Tzu Hsu, Hsu-Hsiang Liu, Yi Kung, Chieh-Jou Lin, Chih-Yu Chao

## Abstract

Non-small cell lung cancer (NSCLC), as the predominant subtype of lung cancer, presents a considerable clinical challenge due to its high rates of recurrence and the significant adverse effects associated with conventional therapeutic modalities. In response to these challenges, this study explored the new combined anticancer effects of epigallocatechin gallate (EGCG) together with thermal cycling-stimulation (TCS). The findings demonstrated that the combination of EGCG and TCS synergistically decreased the viability of A549 and NCI-H460 NSCLC cells, while exhibiting minimal cytotoxic effects on IMR-90 normal lung fibroblasts. Further investigation revealed that EGCG mitigated the TCS-induced upregulation of heat shock proteins HSP70 and HSP105 and concurrently diminished the expression levels of proteasome subunits. This combined effect disrupted proteostasis, resulting in pronounced endoplasmic reticulum (ER) stress. Subsequently, a positive feedback mechanism was established between inositol 1,4,5-trisphosphate receptor (IP_3_R)-mediated ER Ca^2+^ release and excessive reactive oxygen species (ROS) production, ultimately leading the cells to undergo mitochondrial apoptosis. This combined treatment reduces the necessary dosage of EGCG, thereby overcoming limitations related to its poor bioavailability and systemic toxicity, while also preventing the development of thermotolerance induced by TCS. Consequently, this method offers a new and potentially practical therapeutic strategy for treating NSCLC.

## Introduction

Lung cancer is the leading cause of cancer-related deaths worldwide, with non-small cell lung cancer (NSCLC) representing about 85% of all cases (1,2). Although there have been improvements in surgery, chemotherapy, and targeted therapies, the prognosis for patients with advanced NSCLC remains poor, primarily due to the development of drug resistance associated with current treatment modalities (3). Therefore, there is an urgent clinical need to develop novel therapeutic approaches that demonstrate both high efficacy and minimal adverse effects.

Hyperthermia, employed as an adjuvant anticancer treatment, works by disrupting intracellular proteostasis through the elevation of temperature within tumor tissues (4). This thermal stress leads to the accumulation of unfolded proteins, thereby inducing endoplasmic reticulum (ER) stress (4). When ER stress is severe or sustained, it can initiate apoptotic pathways via mechanisms including the disturbance of intracellular calcium ion (Ca^2+^) homeostasis and the excessive generation of reactive oxygen species (ROS) (4,5). However, conventional hyperthermia techniques are often limited by the risk of thermal damage to adjacent normal tissues. To overcome this issue, our research group has developed a novel hyperthermic approach termed thermal cycling-stimulation (TCS). Unlike continuous heating, TCS involves alternating cycles of high and low temperatures, which has been demonstrated to minimize damage to normal cells while preserving substantial inhibitory effects on cancer cell viability (6–10). Moreover, our previous research has shown that TCS works synergistically with various anticancer drugs, including herbal compounds and targeted therapies (6–10). This synergy facilitates a reduction in the required drug dosages and enhances selective anticancer efficacy.

Although hyperthermia holds therapeutic potential, its clinical effectiveness is constrained by the activation of cellular protective mechanisms in response to heat stress. Exposure to heat stress quickly triggers the upregulation of heat shock proteins (HSPs) within cancer cells (11). These molecular chaperones assist in the refolding of denatured proteins, thereby preserving proteostasis, which can lead to thermotolerance and diminish the susceptibility of cancer cells to hyperthermia-induced cytotoxicity (11,12). In addition to protein folding, the ubiquitin-proteasome system (UPS) contributes to proteostasis by degrading abnormal proteins, thereby helping cells withstand stressful conditions (13). Hence, molecular inhibitors that target HSPs or UPS have been proposed as promising approaches to potentiate the anticancer efficacy of hyperthermia (11,14,15).

On the other hand, epigallocatechin gallate (EGCG), the most prevalent and biologically active polyphenol found in green tea, is recognized for its potential against NSCLC (16). Importantly, earlier research has indicated that EGCG can suppress the expression of HSP70 and inhibit proteasome activity (17,18). Additionally, EGCG has been shown to induce ROS production, leading to mitochondrial dysfunction in cancer cells (19). Despite these promising effects, the clinical application of EGCG is limited by its poor chemical stability and low bioavailability, which restricts its accumulation in tumor tissues at therapeutically effective levels (20). Consequently, strategies to enhance the efficacy of EGCG are essential for its successful translation into clinical use.

In this study, we investigated the synergistic anticancer effects of EGCG in combination with TCS on A549 NSCLC cells. Building upon evidence that EGCG inhibits HSPs and UPS, our findings demonstrate that the combined EGCG and TCS exacerbates ER stress. Further mechanistic analysis showed that this combined treatment amplifies a positive feedback loop between calcium ion (Ca^2+^) release from the ER via inositol 1,4,5-trisphosphate receptor (IP_3_R) channels and ROS generation, ultimately inducing mitochondrial apoptosis. Importantly, the combination of EGCG and TCS exhibited no cytotoxic effects on IMR-90 normal lung fibroblast cells, underscoring the safety of this therapeutic approach. Overall, this study aims to elucidate the synergistic anticancer effects of EGCG and TCS and their underlying molecular mechanisms, offering a promising strategy to overcome the limitations associated with heat-induced cytoprotective responses and the inherent instability and poor bioavailability of EGCG. These findings suggest a safe and potentially viable clinical intervention for NSCLC treatment.

## Materials and methods

### Cell culture

Human NSCLC cell lines A549 (cat. no. 60074) and NCI-H460 (cat. no. 60373), as well as the human normal lung fibroblast cell line IMR-90 (cat no. 60204), were obtained from the Bioresource Collection and Research Center of the Food Industry Research and Development Institute. Both A549 and NCI-H460 cells were cultured in Dulbecco’s Modified Eagle Medium (HyClone; Cytiva), whereas IMR-90 cells were maintained in Eagle’s Minimum Essential Medium (Corning, Inc.). All culture media were supplemented with 10% fetal bovine serum (HyClone; Cytiva) and 1% (v/v) penicillin-streptomycin (Gibco; Thermo Fisher Scientific, Inc.). All cell lines were maintained at 37°C in a humidified incubator with 5% CO_2_. Cells were subcultured or harvested for subsequent experiments by 0.05% trypsin-0.5 mM EDTA solution (Gibco; Thermo Fisher Scientific, Inc.).

### The TCS exposure

The TCS was performed using a modified PCR system equipped with waterproof capabilities in the heating area, as described in previous studies (6–8). A minimal volume of water was employed as a thermal conduction medium, enabling direct placement of cell culture dishes onto the heating area. The modified PCR apparatus controlled both heating and cooling phases. The TCS exposure comprised 15 cycles, each consisting of a 3-min high-temperature phase followed by a 30-sec passive cooling interval (Fig. 1A). This heating-cooling period was established based on prior investigations involving various cancer cell lines (6,7). The low-temperature setting was maintained at 37°C to simulate physiological human body temperature. Actual temperatures experienced by cells at the well bottom were monitored using a needle thermocouple. Temperature cycling reached a steady state after the third cycle, with the recorded temperature cycling between 41.4°C and 43.8°C, as depicted in Fig. 1B.

**Figure 1.**
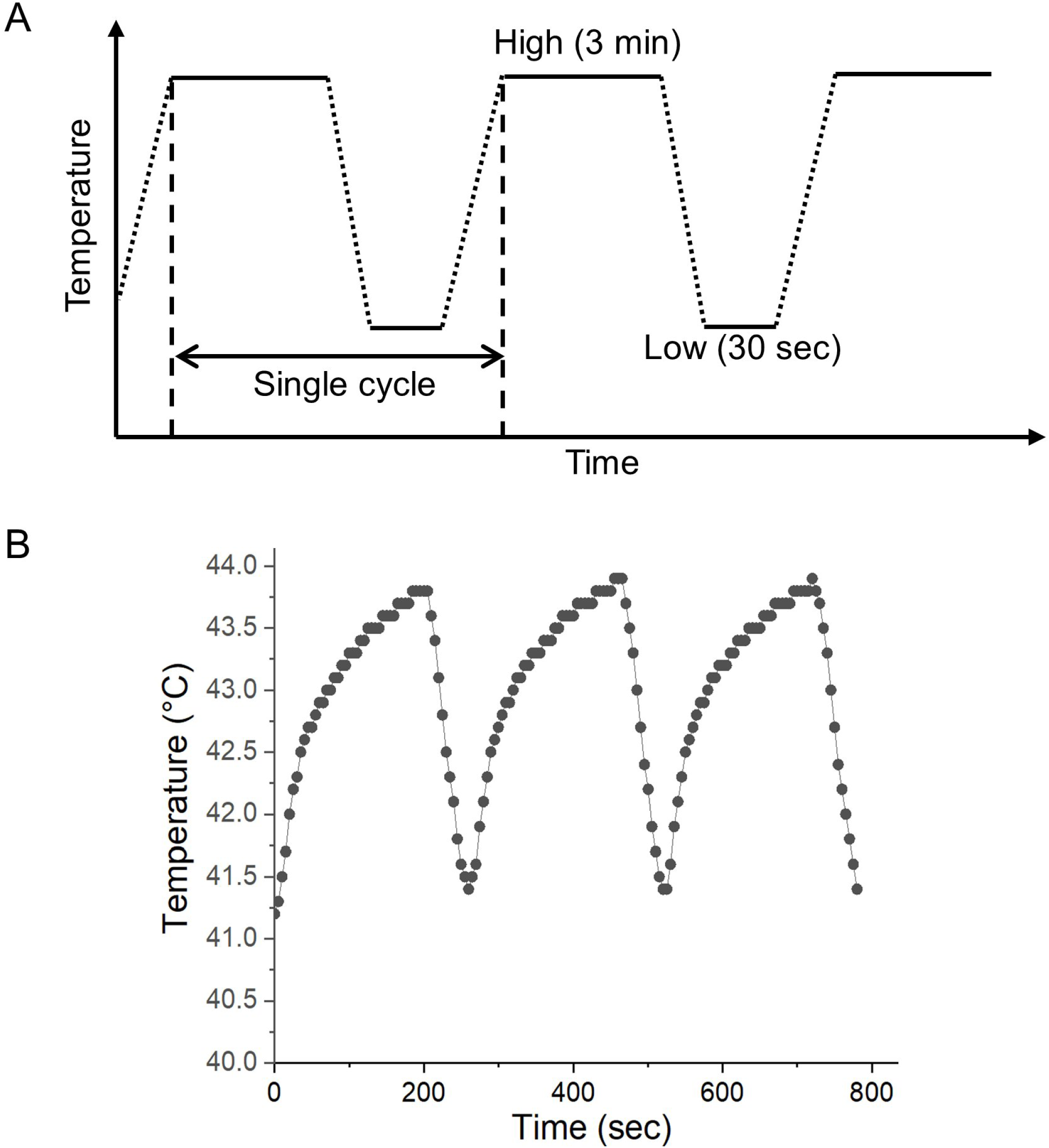
In vitro application of TCS. (A) Schematic diagram of the TCS parameter settings in the modified PCR system. (B) Real-time temperature variation at the bottom of the well, measured with a needle thermocouple. Temperature readings were taken every 5 sec during heating applications.

### Cell treatments

EGCG (MedChemExpress), 2-APB (MedChemExpress), and N-acetylcysteine (NAC, Sigma-Aldrich; Merck KGaA) were dissolved in sterile distilled water to prepare stock solutions at concentrations of 20 mM, 5 mM, and 500 mM, respectively. On the other hand, GSK2606414 (MedChemExpress), JNJ-28583113 (MedChemExpress), and SB-705498 (Sigma-Aldrich; Merck KGaA) were solubilized in dimethyl sulfoxide (DMSO, Sigma-Aldrich; Merck KGaA) at concentrations of 1 mM, 5 mM, and 5 mM, respectively, to make stock solutions. All stock solutions were stored at −20°C until further use. Cells were seeded in 96-well plates at 3,000 cells/well or in 6-well plates at 100,000 cells/well and incubated overnight at 37°C to facilitate cell adherence. For the combined treatment of EGCG and TCS, EGCG was administered immediately prior to TCS exposure. Cells were pretreated with NAC or molecular inhibitors 1 h before EGCG application. Unless specifically indicated, cells were maintained at 37°C in a humidified incubator for an additional 24 h after treatment for further analysis.

### Cell viability assay

Cell viability was assessed using the 3-(4,5-dimethylthiazol-2-yl)-2,5-diphenyltetrazolium bromide (MTT) (Sigma-Aldrich; Merck KGaA) assay. MTT powder was dissolved in distilled water to make a 5 mg/ml stock solution, which was stored at 4°C. Immediately prior to use, the MTT stock solution was diluted with culture medium to a final concentration of 0.5 mg/ml. At 24 h after the treatment, the culture medium in the 96-well plates was replaced with 100 µl of the MTT working solution. The cells were then incubated at 37°C in the dark for 4 h, during which mitochondrial dehydrogenases in viable cells reduced MTT to insoluble formazan crystals. To dissolve the formazan crystals, 100 µl of 10% sodium dodecyl sulfate (SDS) (Bioshop Canada Inc.) solution in 0.01 N hydrochloric acid (HCl) (Echo Chemical Co. Ltd.) was added to each well. After overnight incubation, the absorbance was measured at 570 nm with background subtraction at 690 nm using a Multiskan GO spectrophotometer (Thermo Scientific). Cell viability was expressed as a ratio based on the relative quantity of formazan compared to the untreated control.

### Calculation of synergy quotient (SQ)

The SQ value was calculated by dividing the net inhibitory effect on cell viability from the combined treatment [EGCG + TCS] by the sum of the net inhibitory effects of the individual treatments [EGCG] and [TCS], as expressed by the formula: SQ = [EGCG + TCS] / {[EGCG] + [TCS]} (6).

### Western blot analysis

Protein levels in A549 cells were assessed using Western blotting. After treatment, cells were washed with phosphate-buffered saline (PBS) and then lysed on ice for 1 h using RIPA lysis buffer (50 mM Tris-HCl, pH 7.4, 0.15 M NaCl, 0.25% deoxycholic acid, 1% NP-40, 1% Triton X-100, 0.1% SDS, 1 mM EDTA) (Millipore) supplemented with protease and phosphatase inhibitor cocktails (Millipore). Lysates were centrifuged at 10,000 × g for 30 min at 4°C, and the supernatants were then collected. Protein concentrations were determined using the Bradford protein assay (Bioshop Canada Inc.). Equal amounts of protein samples (20 μg) were separated by sodium dodecyl sulfate polyacrylamide gel electrophoresis (12% acrylamide gel) and then transferred onto polyvinylidene fluoride membranes (Millipore). Membranes were blocked with 5% bovine serum albumin (BSA) in TBST (20 mM Tris, 150 mM NaCl, and 0.1% Tween 20) for 1 h at room temperature, followed by overnight incubation at 4°C with primary antibodies targeting PSMA3, PSMC3 (Cell Signaling Technology, Inc.), HSP70, HSP105, phospho-eIF2α (p-eIF2α; Ser51), total-eIF2α (t-eIF2α), and GAPDH (GeneTex, Inc.). After washing with TBST, the membranes were incubated overnight at 4°C with horseradish peroxidase-conjugated secondary antibodies (Jackson ImmunoResearch Laboratories, Inc.). Antibody dilutions were prepared according to manufacturer’s recommended optimal concentrations. Finally, the immunoreactive signals were visualized using an enhanced chemiluminescence substrate (Advansta) and detected in an Amersham Imager 600 imaging system (GE Healthcare Life Sciences). Protein bands were quantified using Image Lab software (Bio-Rad Laboratories, Inc.).

### Measurement of cytosolic Ca^2+^ levels

Cytosolic Ca^2+^ levels were measured using the fluorescent dye Fluo-4 AM (MedChemExpress) to quantify cytosolic Ca^2+^ levels. A 1 mM stock solution of Fluo-4 AM was prepared by dissolving the dye in DMSO and stored at −20°C until further use. At 24 h after treatments, cells were harvested and washed with Hanks’ Balanced Salt Solution (HBSS), and then incubated with 1 μM Fluo-4 AM in HBSS for 1 h at 37°C in the dark to load the dye. Following the incubation, cells were washed with HBSS and finally resuspended in fresh HBSS. Fluorescence intensity of Fluo-4 AM was measured using a FACSLyric flow cytometer (BD Biosciences) in the FITC channel. Cytosolic Ca^2+^ levels were quantified by calculating the mean fluorescence intensity and expressed as fold changes relative to the untreated control group.

### Measurement of intracellular ROS levels

The dihydroethidium (DHE) (Sigma-Aldrich; Merck KGaA) fluorescent dye was employed to measure ROS levels in A549 cells. At 24 h after treatments, cells were harvested and washed with PBS prior to staining. Cells were then incubated with a 5 μM DHE working solution for 30 min at 37°C in the dark. Fluorescence intensity was measured using a FACSLyric flow cytometer (BD Biosciences) in the PE channel. ROS levels were determined by calculating the mean fluorescence intensity and expressed as fold changes compared to the control group.

### Mitochondrial membrane potential (MMP) analysis

The MMP integrity was assessed using the JC-1 (MedChemExpress) fluorescent dye. At 24 h after treatments, cells were harvested and rinsed with PBS. Subsequently, cells were incubated with 5 μg/ml JC-1 in PBS for 30 min at 37°C in the dark. Fluorescence intensities of JC-1 were measured using the FACSLyric system (BD Biosciences) in both the PE (red fluorescence) and FITC (green fluorescence) channels. MMP disruption was quantified by the proportion of cells exhibiting a fluorescence shift from red (JC-1 aggregates) to green (JC-1 monomers).

### Apoptosis analysis

Apoptotic rates were quantified using an Annexin V-FITC and propidium iodide (PI) double staining kit (BD Biosciences). At 24 h after treatments, cells were collected, washed with PBS, and then incubated in a binding buffer containing Annexin V-FITC and PI for 30 min at 37°C in the dark, according to the manufacturer’s instructions. Fluorescence intensities were measured using the FACSLyric system (BD Biosciences) in both the PE and FITC channels. Early and late apoptotic cells were quantified as the percentage of cells identified as (PE-/FITC+) and (PE+/FITC+), respectively. The percentages of these populations were calculated accordingly.

### Statistical analysis

All experiments were performed in triplicate, and data are presented as mean ± standard deviation. Statistical analyses were conducted using one-way analysis of variance (ANOVA), followed by Tukey’s post hoc test, using the OriginPro 2015 software (version 92E, OriginLab Corporation). A *p*-value less than 0.05 was considered statistically significant. For clarity, only selected pairwise comparisons are denoted in the figures.

## Results

### Synergistic and selective inhibitory effects of EGCG and TCS on NSCLC cell viability

To assess the effect of EGCG and TCS on the viability of A549 cells, the MTT assay was performed 24 h after the treatment. As illustrated in Fig. 2A, EGCG induced a significant, dose-dependent reduction in A549 cell viability. On the other hand, TCS alone caused only a slight decrease in A549 cell viability, maintaining about 90% of control levels. Notably, the combined administration of EGCG and TCS produced a more substantial inhibition effect on A549 cell viability. The SQ values calculated in Table 1A confirmed a synergistic cytotoxic effect of EGCG and TCS, with SQ values exceeding 1 at EGCG concentrations ranging from 30 µM to 60 µM. Consistent with the MTT results, morphological evaluation using a light microscope at 24 h post-treatment (Fig. 2B) demonstrated that the combined treatment of EGCG and TCS induced pronounced morphological alterations in A549 cells, including reduced cell density, cellular shrinkage, and an increased proportion of non-adherent cells. To further validate that this effect is widely applicable, the MTT assay was also performed on a different NSCLC cell line, NCI-H460. Results shown in Fig. 2C confirmed a synergistic cytotoxic effect of EGCG and TCS in NCI-H460 cells, with corresponding SQ values presented in Table 1B. Combining the data from Tables 1A and 1B, the combination of 50 µM EGCG and TCS produced SQ values greater than 2 in both A549 and NCI-H460 cell lines, thereby justifying the selection of 50 µM EGCG for subsequent experiments in this study. Notably, as shown in Fig. 2D, the combination of EGCG and TCS did not cause a reduction in the viability of IMR-90 normal lung fibroblasts, indicating that this combined treatment is relatively safe. Overall, these results show that EGCG and TCS work synergistically to inhibit the viability of NSCLC cell lines A549 and NCI-H460 while having no observed adverse effects on IMR-90 normal lung fibroblast cells, thereby indicating a selective anticancer effect.

**Figure 2.**
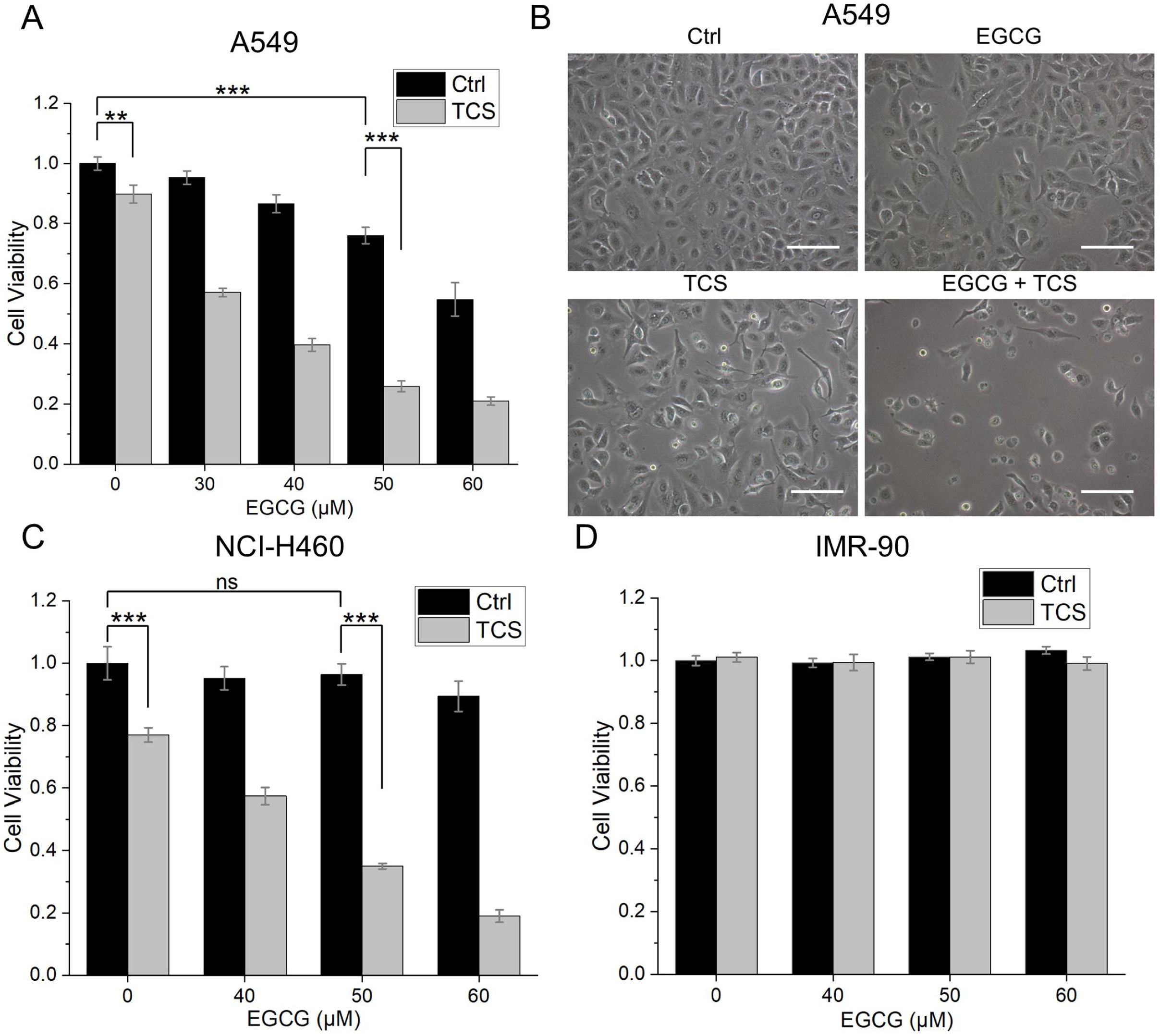
Effects of TCS and EGCG alone or their combination treatment on the viability and morphology of A549, NCI-H460, and IMR-90 cells. (A) The viability of A549 cells was evaluated using the MTT assay 24 h after treatment with various concentrations of EGCG alone or in combination with TCS. (B) Cellular morphology was examined through light microscopy 24 h after treatment with 50 µM EGCG and/or TCS. Scale bar = 100 μm. (C) The viability of NCI-H460 cells was measured 24 h after treatment with different concentrations of EGCG alone or combined with TCS. (D) The viability of IMR-90 cells was similarly assessed 24 h after treatment with various concentrations of EGCG alone or in combination with TCS. Statistical significance between the indicated groups is denoted as ** *p* < 0.01 and *** *p* < 0.001, while non-significant differences are labeled as ns.

**Table 1.**
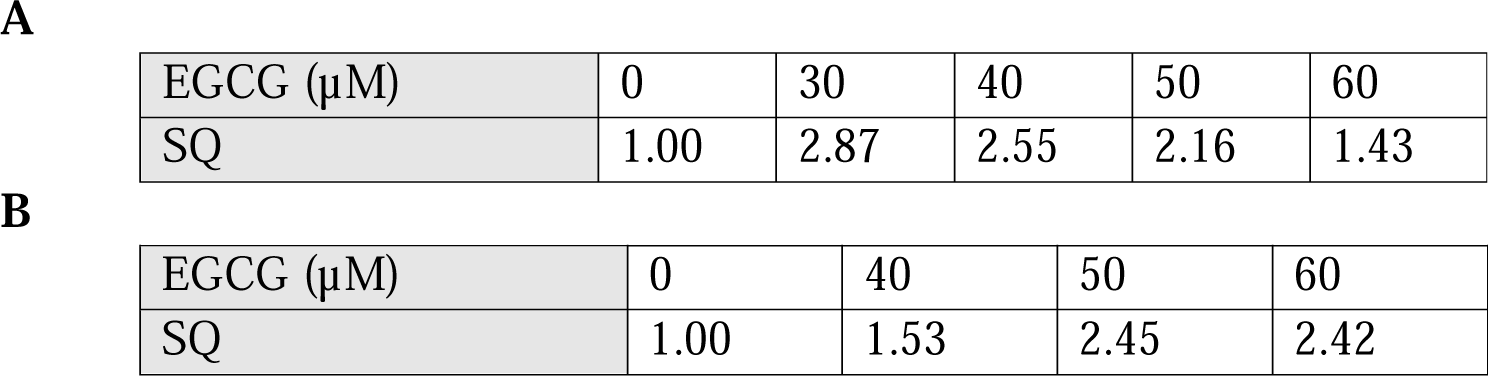
Synergy quotient (SQ) for EGCG in combination with TCS in (A) A549 and (B) NCI-H460 cells.

### Effects of EGCG and TCS on the expression levels of HSPs and proteasome subunits in A549 cells

To understand the mechanism underlying the synergistic effect of EGCG and TCS, we first investigated whether these treatments influence the regulation of HSPs in A549 cells. Western blot analysis was employed to quantify the expression levels of HSP70 and HSP105. As shown in Fig. 3A and 3B, TCS treatment increased the expression of both HSP70 and HSP105 in A549 cells, whereas co-treatment with EGCG significantly reduced this induction. In addition to HSPs, the expression levels of proteasome subunits PSMA3 and PSMC3 were also evaluated by Western blotting. Fig. 3C and 3D shows that TCS alone exerted a negligible effect on PSMA3 expression and induced only a slight reduction in PSMC3 levels. In contrast, EGCG was found to significantly decrease the expression of both PSMA3 and PSMC3 in A549 cells treated with TCS. Collectively, these results suggest that EGCG may potentiate the sensitivity of A549 cells to TCS by downregulating HSP70, HSP105, and proteasome subunits, thereby causing the accumulation of unfolded proteins.

**Figure 3.**
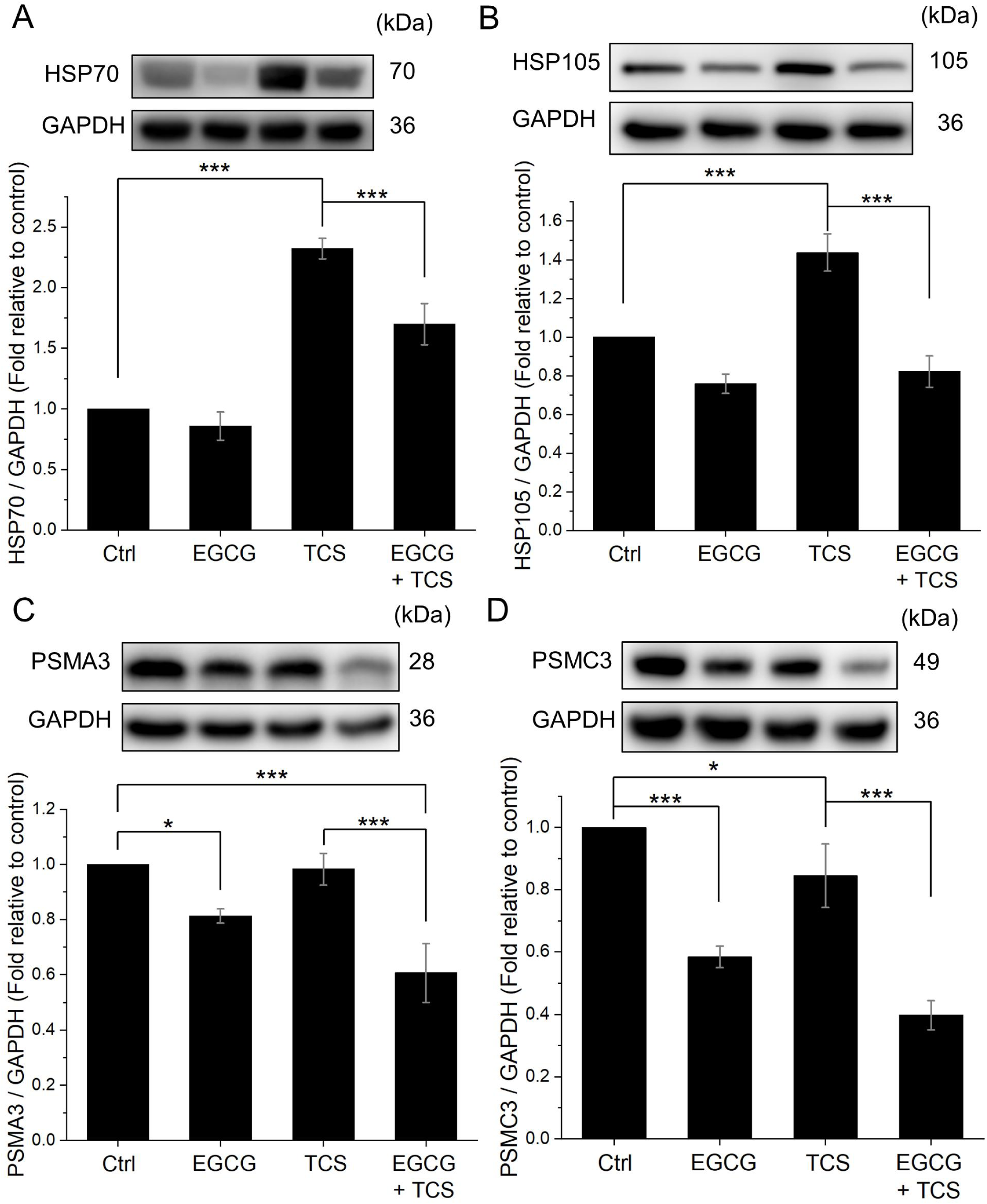
Effects of TCS, EGCG, and their combined application on the expression levels of HSPs and proteasome subunits in A549 cells. Western blot analysis was employed to measure the expression levels of (A) HSP70, (B) HSP105, (C) PSMA3, and (D) PSMC3 were examined by Western blot analysis. GAPDH was used as the internal loading control. Statistical significance between the indicated groups is indicated as * *p* < 0.05 and *** *p* < 0.001.

### EGCG and TCS induce ER stress in A549 cells

The intracellular accumulation of unfolded proteins is known to provoke ER stress, which, if sustained or severe, can culminate in cellular damage (21). This study further explored the involvement of the PERK-eIF2α signaling axis, a key ER stress pathway, in mediating cell death induced by EGCG and TCS. As shown in Fig. 4A, administration of the PERK inhibitor GSK2606414 partially mitigated the reduction in A549 cell viability caused by combined EGCG and TCS treatment in a dose-dependent manner. Additionally, Western blot analysis of downstream signaling revealed that TCS significantly enhanced the phosphorylation of eIF2α, with the combined treatment of EGCG and TCS producing an even more pronounced effect (Fig. 4B). These findings indicate that the combination of EGCG and TCS effectively induces ER stress, which partially contributes to the observed decrease in viability of A549 cells.

**Figure 4.**
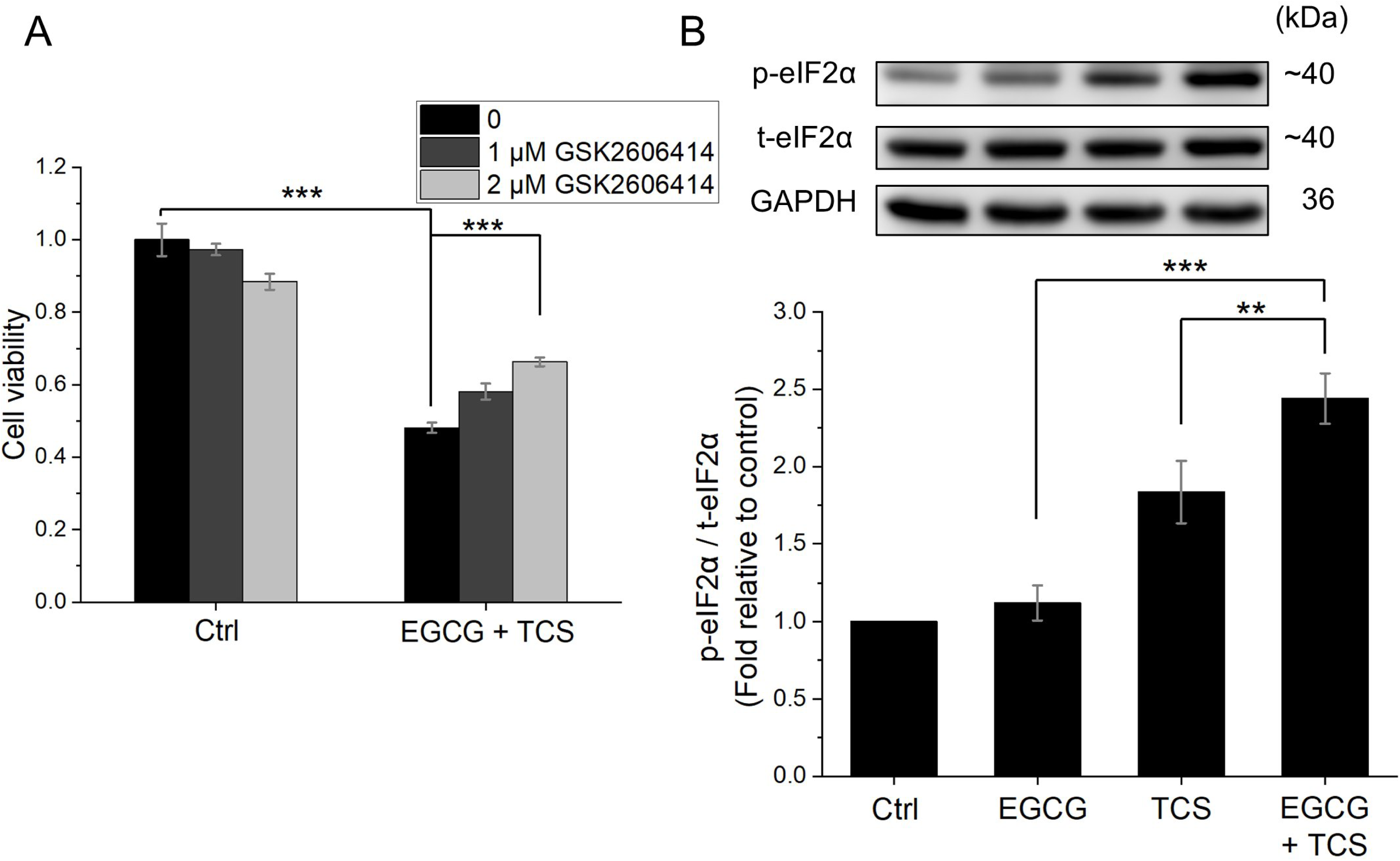
The effects of TCS and EGCG on the PERK-eIF2α signaling pathway in A549 cells were evaluated. (A) Cell viability of A549 cells was measured using the MTT assay 24 h following combined treatment with EGCG and TCS, in the presence or absence of the PERK inhibitor GSK2606414, which was administered 1 h prior to the combined treatment. (B) The protein levels of p-eIF2α and t-eIF2α were examined by Western blot analysis 3 h after the treatment. GAPDH was utilized as the internal loading control. Statistical significance between the indicated groups is indicated as ** *p* < 0.01 and *** *p* < 0.001.

### Induction of IP_3_R-mediated ER Ca^2+^ release by the combined treatment of EGCG and TCS in A549 cells

Under ER stress, Ca^2+^ stored in the ER can be released via IP_3_R channels, potentially leading to intracellular Ca^2+^ dysregulation and subsequent cellular damage (22). Based on our previous findings in Fig. 4, cytosolic Ca^2+^ levels were further quantitatively measured using the Fluo-4 AM fluorescent dye through flow cytometric analysis. As shown in Fig. 5A and 5B, treatment with EGCG alone did not significantly affect cytosolic Ca^2+^ levels in A549 cells, whereas TCS alone caused only a slight increase. In contrast, the combination of EGCG and TCS produced a substantial elevation in cytosolic Ca^2+^ levels. Notably, this increase was greatly attenuated by the IP_3_R inhibitor 2-APB, indicating that the elevation in Ca^2+^ level is mediated by the activation of IP_3_R channels. To further elucidate the role of ER Ca^2+^ release in the synergistic cytotoxicity induced by EGCG and TCS in A549 cells, a rescue experiment employing 2-APB was conducted using the MTT viability assay. As shown in Fig. 5C, 2-APB significantly lessened the combined treatment’s reduction of A549 cell viability in a dose-dependent manner. In addition to IP_3_R channels, 2-APB is also known to affect thermosensitive TRP channels such as TRPV1 and TRPM2 (23,24). However, we found that selective inhibitors for TRPM2 (JNJ-28583113) and TRPV1 (SB-705498) did not reverse the synergistic cytotoxic effects of EGCG and TCS (Fig. 5D), suggesting that these TRP channels are unlikely to play a significant role in this process. Taken together, these findings confirm the essential role of IP_3_R channels and the resultant ER Ca^2+^ release in the synergistic anticancer effects induced by the combined EGCG and TCS treatment.

**Figure 5.**
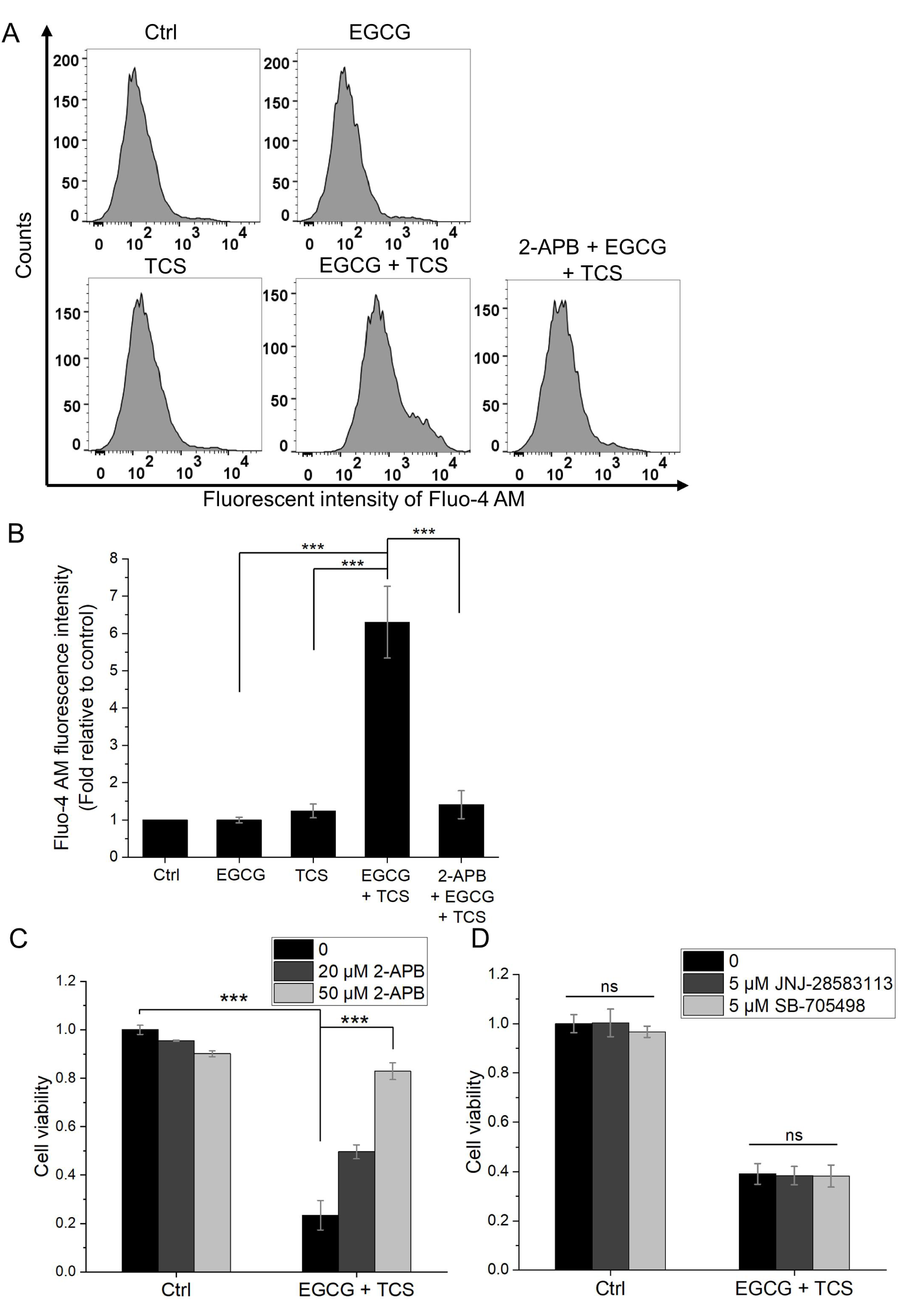
Investigation of the effects of TCS and EGCG on the Ca^2+^ homeostasis in A549 cells. (A) Cytosolic Ca^2+^ levels in A549 cells were measured in A549 cells utilizing the fluorescent dye Fluo-4 AM via flow cytometry. (B) Quantitative analysis of the mean fluorescence intensities of Fluo-4 AM as shown in panel (A). (C) Cell viability of A549 cells was evaluated by the MTT assay 24 h following combined EGCG and TCS treatment, in the presence or absence of the IP_3_R inhibitor 2-APB. The 2-APB inhibitor was administered 1 h prior to the combined EGCG and TCS treatment. (D) Assessment of A549 cell viability was also performed by the MTT assay 24 h after combined EGCG and TCS treatment, with or without the TRPM2 inhibitor JNJ-28583113 and the TRPV1 inhibitor SB-705498. Similarly, these inhibitors were applied 1 h before the EGCG and TCS treatment. Statistical significance between the indicated groups is denoted as *** *p* < 0.001, while non-significant differences are labeled as ns.

### The combination of EGCG and TCS induces ROS production in A549 cells

ER stress and intracellular Ca^2+^ dysregulation can cause excessive ROS generation (5). In this study, intracellular ROS levels in A549 cells were measured using the fluorescent dye DHE via flow cytometry. As shown in Fig. 6A and the quantification data in Fig. 6B, treatment with either EGCG or TCS alone resulted in only a slight and statistically non-significant increase in ROS levels. Conversely, the combined administration of EGCG and TCS produced a pronounced and statistically significant elevation of ROS levels in A549 cells. To explore the role of ROS in the reduction of cell viability caused by this combination, a rescue experiment employing the ROS scavenger NAC was conducted using the MTT assay. The results shown in Fig. 6C indicate that NAC effectively mitigated the synergistic cytotoxicity induced by the EGCG and TCS combination in A549 cells, suggesting that the excessive ROS production induced by the combined treatment is essential for the synergistic cytotoxicity.

**Figure 6.**
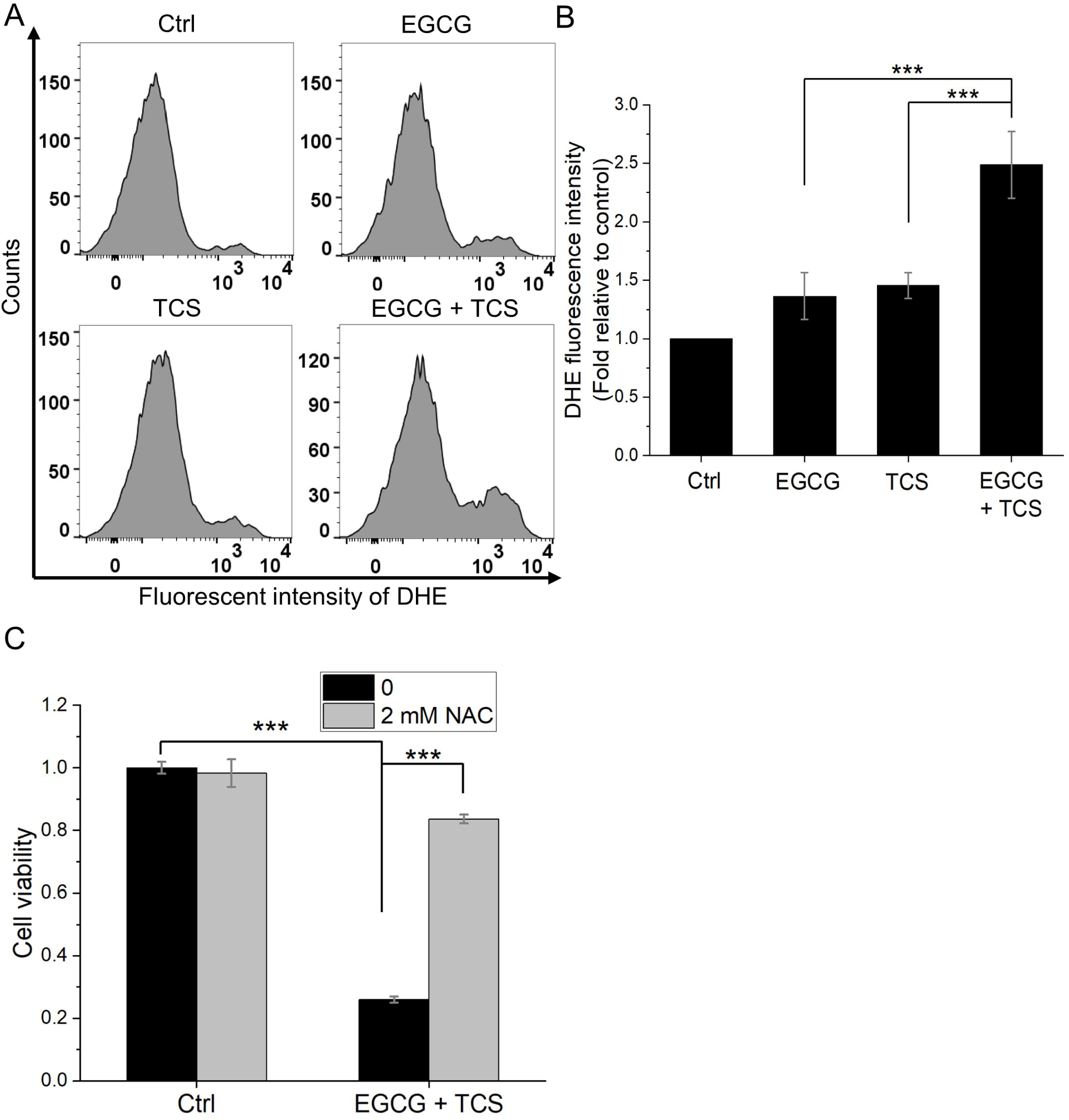
Evaluation of the effects of TCS and EGCG on ROS regulation in A549 cells. (A) Intracellular ROS levels in A549 cells were determined using the fluorescent dye DHE via flow cytometry. (B) Quantification of mean fluorescent intensities of DHE as shown in panel (A). (C) A549 cell viability was measured by the MTT assay 24 h following combined treatment with EGCG and TCS, with or without the ROS scavenger NAC. NAC was administered 1 h prior to the combined treatment. Statistical significance between the indicated groups is denoted as *** *p* < 0.001.

### Mutual amplification of ROS production and ER Ca^2+^ release under combined EGCG and TCS treatment in A549 cells

To explore the mechanistic relationship between ER Ca^2+^ release and ROS generation, ROS levels in A549 cells were first assessed by DHE staining in the presence of 2-APB, an inhibitor of IP_3_R-mediated Ca^2+^ release. As shown in Fig. 7A and 7B, 2-APB significantly attenuated the ROS elevation triggered by the combined EGCG and TCS treatment, indicating that ER Ca^2+^ release through IP_3_R channels is essential for the substantial increase in ROS levels. On the other hand, the cytosolic Ca^2+^ levels were measured using Fluo-4 AM staining in the presence of NAC (Fig. 7C and 7D). We found that NAC treatment effectively suppressed the rise in cytosolic Ca^2+^ levels induced by the combined treatment, which suggests that elevated ROS levels contribute to ER Ca^2+^ release. Collectively, these data support the existence of a positive feedback loop between the ER Ca^2+^ release and ROS generation, wherein mutual amplification of these signals culminates in enhanced cell death.

**Figure 7.**
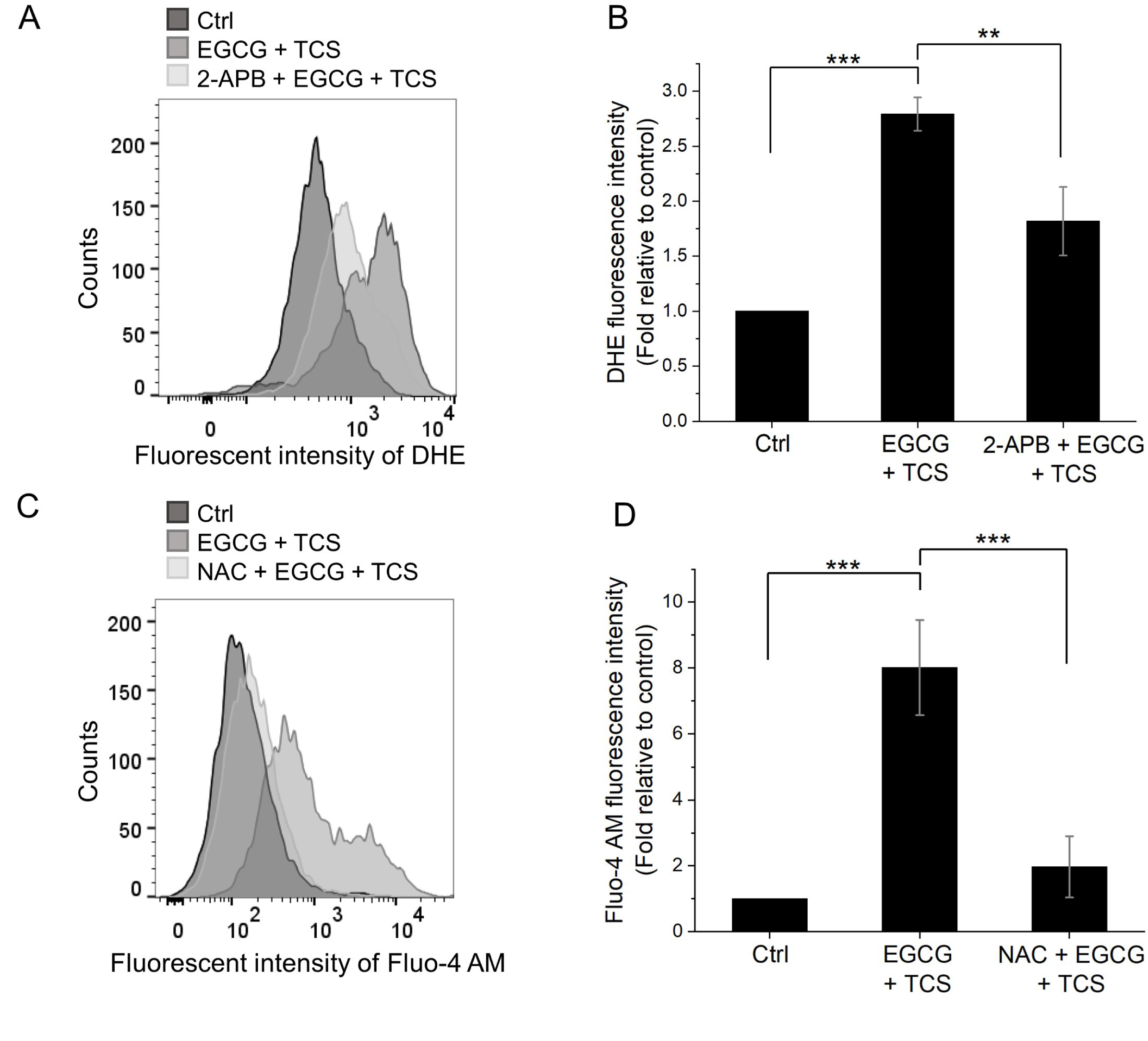
Effects of TCS and EGCG on the ROS-Ca^2+^ feedback loop in A549 cells. (A) Intracellular ROS levels in A549 cells were measured using the fluorescent dye DHE via flow cytometry, following combined treatment with EGCG and TCS in the absence or presence of 2-APB. (B) Quantitative analysis of the mean fluorescent intensities of DHE shown in panel (A). (C) Cytosolic Ca^2+^ levels in A549 cells were assessed using the fluorescent dye Fluo-4 AM via flow cytometry following combined EGCG and TCS treatment, in the absence or presence of NAC. (D) Quantification of the mean fluorescent intensities of Fluo-4 AM presented in panel (C). Statistical significance between the indicated groups is indicated as ** *p* < 0.01 and *** *p* < 0.001.

### The combination of EGCG and TCS induces mitochondrial apoptosis in A549 cells

Excessive ROS accumulation and mitochondrial Ca^2+^ overload are known to cause mitochondrial damage (25,26). Given that the combined EGCG and TCS treatment effectively triggered ER Ca^2+^ release and increased ROS levels, we further evaluated changes in the MMP using the fluorescent dye JC-1 via flow cytometry. As shown in Fig. 8A, the combined EGCG and TCS treatment caused a significant shift in JC-1 fluorescence from red to green, indicative of MMP disruption. Quantitative analysis in Fig. 8B shows that the combined treatment caused a more substantial loss of MMP compared to either treatment alone. Mitochondrial damage may eventually activate the intrinsic apoptotic pathway (25,26). Accordingly, the evaluation of apoptosis was performed using Annexin V-FITC/PI double staining by flow cytometry analysis. As shown in Fig. 8C and 8D, the EGCG and TCS combination significantly increased the proportion of apoptotic cells compared to single treatments, highlighting the strong anticancer potential of this combined treatment.

**Figure 8.**
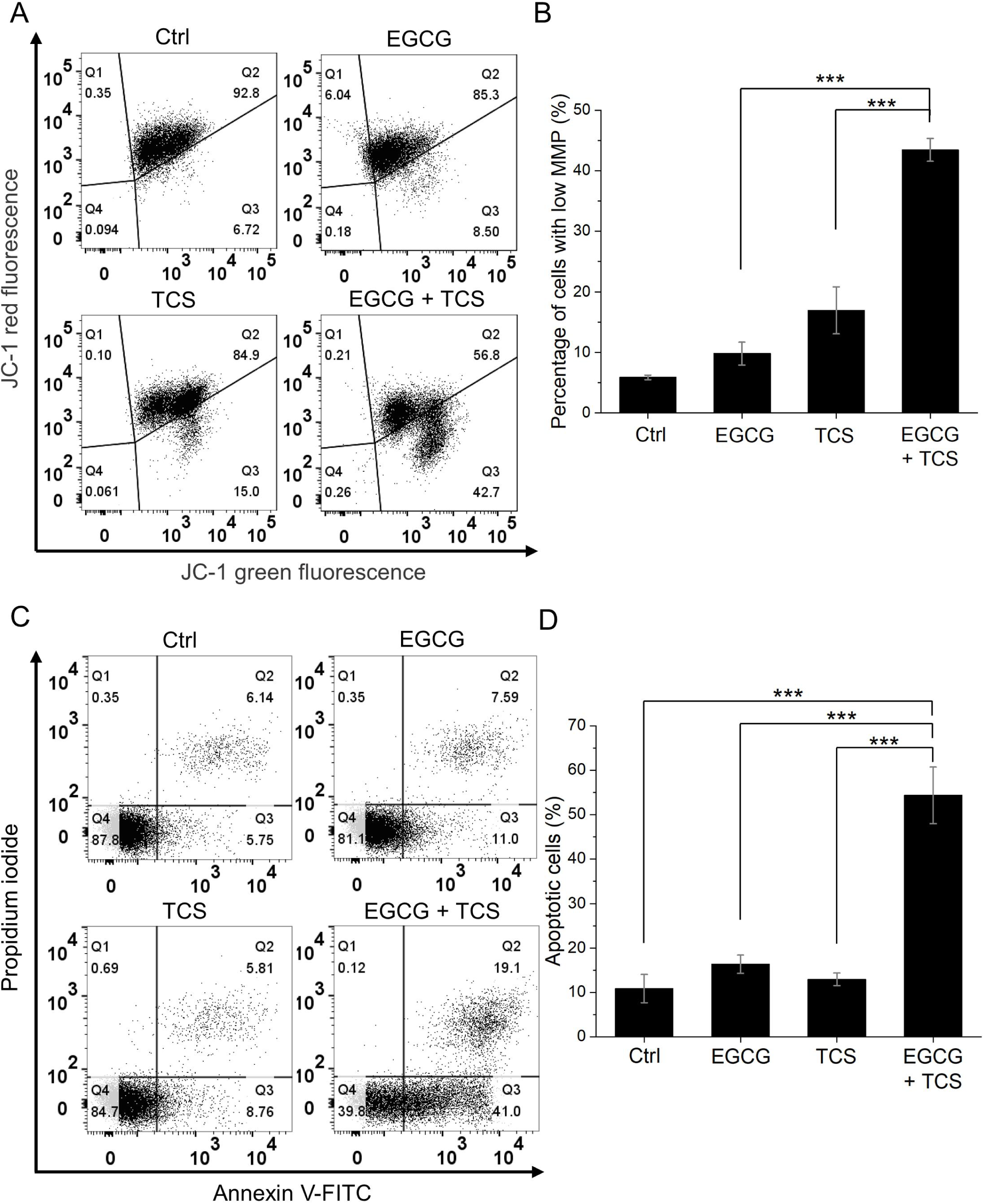
Effects of TCS and EGCG on mitochondrial apoptosis in A549 cells. (A) MMP in A549 cells was evaluated using the fluorescent dye JC-1 by flow cytometry. (B) Quantification of the percentage of cells with low MMP (Q3 regions in panel (A)). (C) Apoptotic A549 cells were detected using the Annexin V-FITC/PI double staining and analyzed via flow cytometry. (D) Quantitative measurement of the percentage of apoptotic cells (Q2 + Q3 regions in panel (C)). Statistical significance between the indicated groups is denoted as *** *p* < 0.001.

## Discussion

The considerable challenges associated with the treatment of NSCLC have prompted extensive investigation into more effective combination therapies (2,6). EGCG, a natural compound, shows strong anticancer properties, but its clinical application has been limited due to poor bioavailability, low stability, and potential hepatotoxicity at elevated doses (20,27). Our data, as illustrated in Fig. 2, indicate that the combined administration of 50 μM EGCG and TCS synergistically suppresses the viability of NSCLC cell lines A549 and NCI-H460, while exhibiting negligible cytotoxic effects on IMR-90 normal lung fibroblasts. This synergistic approach effectively reduces the necessary therapeutic dosage of EGCG, thereby offering a promising strategy to overcome its clinical limitations.

To understand the mechanisms underlying this synergistic effect, we first studied the influence of EGCG and TCS on HSPs and UPS, two critical regulators of intracellular proteostasis. HSPs, particularly HSP70, are often overexpressed in various malignancies including NSCLC, where they facilitate cell proliferation, inhibit apoptosis, and contribute to drug resistance (28–30). Consequently, several molecular inhibitors targeting HSPs have been developed as anticancer agents (11,12). Nonetheless, hyperthermia-induced upregulation of HSP70 and HSP105 is associated with the development of thermotolerance, which diminishes the clinical efficacy of thermal therapies (11,31). Notably, EGCG has been shown to suppress HSP70 expression and competitively bind to its ATPase domain, inhibiting its chaperone function (17). Consistent with these findings, our results demonstrate that EGCG effectively attenuates the TCS-induced elevation of HSP70 and HSP105 levels (Fig. 3A and 3B), suggesting that EGCG may disrupt adaptive thermotolerance and sensitize A549 cells to TCS-induced proteotoxic stress. In addition to HSPs, the UPS plays a crucial role in oncogenesis and progression, rendering it a viable therapeutic target in cancer treatments (15,32). Earlier research has demonstrated that EGCG acts as a blocker to the chymotrypsin-like function of the 20S proteasome (33). Our findings further reveal that the combination of EGCG and TCS markedly decreases the protein expression of proteasome subunits PSMA3 and PSMC3 (Fig. 3C and 3D). Collectively, these results suggest that the synergistic anticancer effects of EGCG and TCS may result from concurrently disrupting HSP-mediated protein folding and UPS-dependent protein degradation. This leads to an accumulation of unfolded proteins, which strongly triggers severe ER stress in NSCLC cells.

ER stress serves as a crucial cellular response to stress and has a dual effect on cancer cells. When cancer cells experience mild proteotoxic stress in the tumor microenvironment, they activate the unfolded protein response (UPR) to enhance survival. However, if ER stress becomes persistent or severe, it overwhelms this adaptive capacity and eventually leads to apoptosis (21,34). Consequently, manipulating this balance to shift the UPR signaling from a pro-survival to a pro-apoptotic mode represents a promising therapeutic strategy for treating NSCLC (34–36). In this study, we demonstrated that pretreatment with the PERK inhibitor GSK2606414 partially restored the reduction in viability of A549 cells caused by the combined treatment of EGCG and TCS (Fig. 4A). Additionally, phosphorylation of the downstream target eIF2α was found to enhance in the combination treatment group (Fig. 4B). These findings indicate that the PERK-eIF2α axis, a key pathway in ER stress, is activated by the combined treatment and contributes to reduced viability in NSCLC cells.

Upon severe ER stress, calcium ions (Ca^2+^) are released from the ER, which can cause further cellular damage (5,22,37). Notably, activation of the PERK pathway has been shown to induce Ca^2+^ release from ER via the IP_3_R (22,37). Consistent with these reports, our data (Fig. 5A and 5B) reveal that the combination of EGCG and TCS significantly elevates cytosolic Ca^2+^ levels. Furthermore, the use of the IP_3_R inhibitor 2-APB notably reduced this Ca^2+^ increase, indicating that the effect is dependent on IP_3_R activity. Concurrently, ER stress is recognized to stimulate ROS production (5,38). Our results in Fig. 6A and 6B confirm that the combined EGCG and TCS treatment effectively increases intracellular ROS levels in A549 cells. Importantly, the release of Ca^2+^ from the ER and the production of ROS are interrelated processes. Previous research has shown that oxidation of key thiol groups in the IP_3_R protein enhances its channel activity (39,40). At the same time, increased cytosolic or mitochondrial Ca^2+^ levels activate ROS-generating enzymes, thereby facilitating free radical formation (41–43). In this study, we found that the increase in cytosolic Ca^2+^ caused by the combination treatment was reduced by the ROS scavenger NAC, while ROS levels were lowered by the IP_3_R inhibitor 2-APB (Fig. 7). These observations suggest a bidirectional crosstalk between Ca^2+^ dysregulation and excessive ROS production. Moreover, treatment with either NAC or 2-APB alone significantly lessened the decrease in cell viability caused by the combined EGCG and TCS treatment in A549 cells (Fig. 5C and 6C), underscoring the essential role of this crosstalk in mediating the synergistic cytotoxic effects observed in A549 cells.

Mitochondrial ROS and Ca^2+^ overload are established triggers for the opening of the mitochondrial permeability transition pore (mPTP), which subsequently leads to the loss of MMP (22,25,26). Building upon our findings that the combination of EGCG and TCS causes intracellular Ca^2+^ dysregulation and ROS generation, we further demonstrated that this combination effectively reduced the MMP in A549 cells, as shown by JC-1 staining (Fig. 8A and 8B). The opening of the mPTP results in mitochondrial matrix swelling and rupture of the outer mitochondrial membrane, facilitating the release of cytochrome c from the intermembrane space into the cytosol, thereby initiating the intrinsic apoptotic pathway (44,45). Consistent with this mechanism, our data in Fig. 8C and 8D reveal a significant increase in the proportion of apoptotic cells following combined EGCG and TCS treatment. Overall, these findings indicate that the combination of EGCG and TCS induces intrinsic apoptosis by promoting mitochondrial ROS accumulation and Ca^2+^ overload in A549 cells.

In summary, this study elucidates the synergistic effects of EGCG and TCS in reducing the viability of NSCLC cells and clarifies the underlying molecular mechanism (Fig. 9). Initially, TCS causes protein unfolding, while EGCG further suppresses the activity and expression of HSPs (e.g., HSP70 and HSP105) and proteasomes, thereby leading to ER stress. This heightened ER stress establishes a positive feedback loop between IP_3_R-mediated ER Ca^2+^ release and excessive ROS production, which ultimately induces mitochondrial apoptosis. Notably, this combination treatment selectively damages NSCLC cells while sparing normal lung cells, potentially attributable to the elevated basal ROS levels and more vulnerable proteostasis regulation in cancer cells (46,47). This combined approach not only reduces the necessary dose of EGCG, overcoming its limited bioavailability and systemic side effects, but also prevents the thermotolerance typically caused by hyperthermia. Further in vivo investigations are warranted to validate these findings and advance the clinical translation of this promising combination therapy.

**Figure 9.**
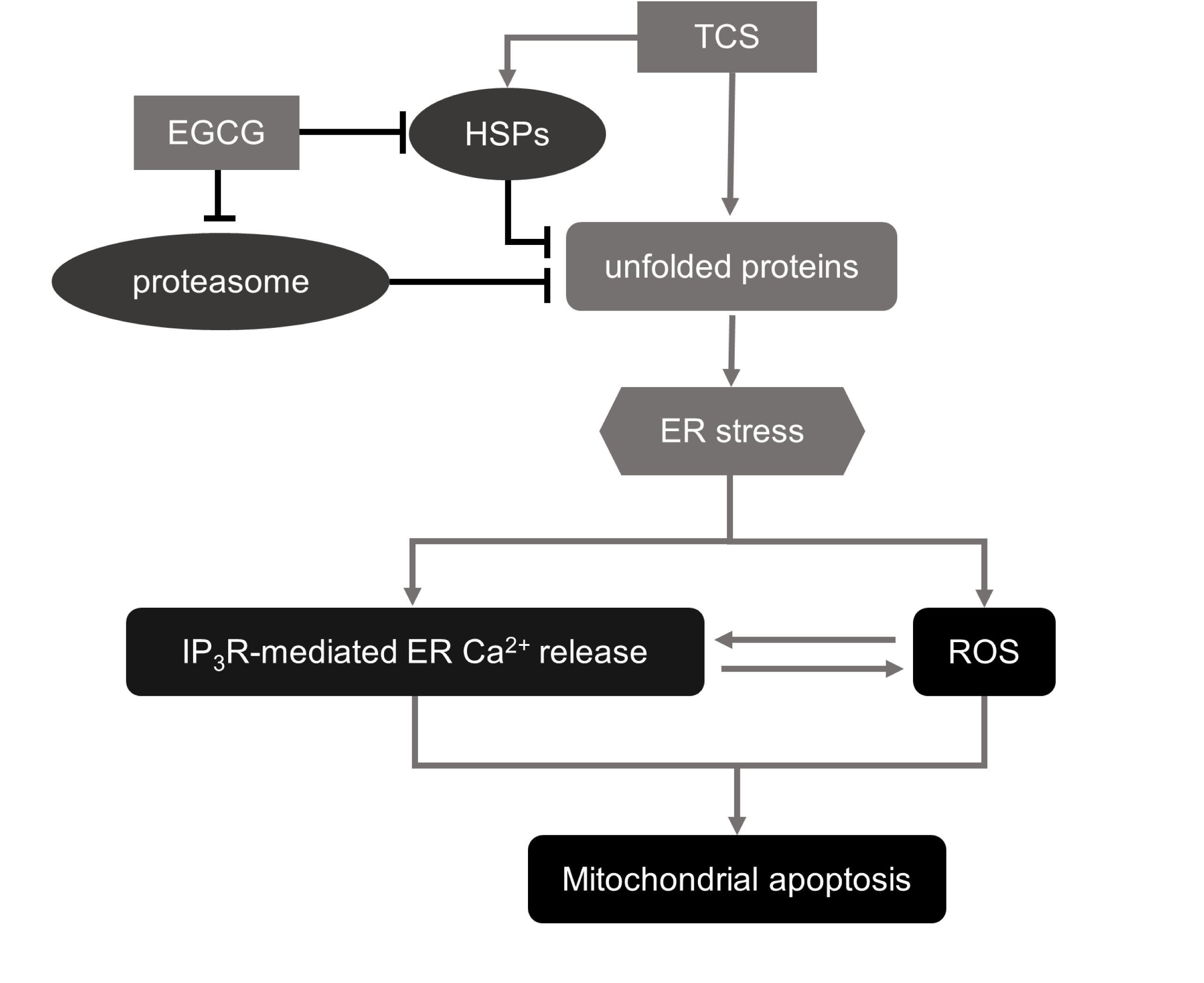
Schematic representation of the proposed molecular mechanisms underlying the synergistic anticancer effects of combined EGCG and TCS treatment in NSCLC cells. TCS induces protein unfolding, whereas EGCG suppresses the activity and expression of HSPs and proteasomes. This combined effect disrupts proteostasis, causing accumulation of unfolded proteins and triggering ER stress. The ER stress then promotes the mutual amplification of ER Ca^2+^ release via IP_3_R and ROS generation, ultimately leading to mitochondrial apoptosis.

## Acknowledgements

The authors would like to acknowledge the service provided by the Research Core Facilities 3 Laboratory of the Department of Medical Research at National Taiwan University hospital for the use of the flow cytometry system.

## Availability of data and materials

The data generated in the present study may be requested from the corresponding author.

## Authors’ contributions

CYC conceived and supervised the study. FTH and CYC designed the study. FTH, HHL, YK, CJL, and CYC confirmed the authenticity of all data. FTH and CYC contributed to data analysis and wrote the manuscript. All authors performed the experiments and approved the final version of the manuscript.

## Ethics approval and consent to participate

Not applicable

## Patient consent for publication

Not applicable

## Abbreviations

NSCLC: non-small cell lung cancer
ER: endoplasmic reticulum
ROS: reactive oxygen species
TCS: thermal cycling-stimulation
HSP: heat shock protein
UPS: ubiquitin-proteasome system
EGCG: epigallocatechin gallate
IP_3_R: inositol 1,4,5-trisphosphate receptor
NAC: N-acetylcysteine
DMSO: dimethyl sulfoxide
MTT: 3-(4,5-dimethylthiazol-2-yl)-2,5-diphenyltetrazolium bromide
SDS: sodium dodecyl sulfate
HCl: hydrochloric acid
SQ: synergy quotient
PBS: phosphate-buffered saline
BSA: bovine serum albumin
p-eIF2α: phospho-eIF2α
t-eIF2α: total-eIF2α
HBSS: Hanks’ Balanced Salt Solution
DHE: dihydroethidium
MMP: mitochondrial membrane potential
FITC: fluorescein isothiocyanate
PI: propidium iodide
ANOVA: one-way analysis of variance
mPTP: mitochondrial permeability transition pore

